# TLR4 mediates Il-6 production in irinotecan-induced mucositis

**DOI:** 10.1101/715417

**Authors:** S. Khan, J. M. Bowen, Hannah R. Wardill

## Abstract

**Introduction:** Irinotecan is a first line chemotherapeutic agent for colorectal carcinoma and well known for debilitating diarrhoea caused by mucositis. Tlr4, a pattern recognition receptor, has been implicated in irinotecan -induced mucositis due to its activation of downstream inflammatory pathways and interaction with luminal microbes. Tlr4 deletion has been reported to attenuate mucosal injury in preclinical models of mucositis, hypothesised to occur by blockade of Il-6 production. As such, the current study aimed to determine the relationship between Tlr4 and Il-6 in the context of irinotecan induced mucositis.

**Materials and methods:** Archived ileum and colon tissues from BALB/c wild type (WT) and Tlr4 -/-billy (Tlr4 KO) female mice were used throughout the study. Mice had received single dose of irinotecan (i.p. 270 mg/kg) or vehicle and sacrificed at 6 and 72 hours. Il-6 mRNA and protein expression was determined by qPCR and immunofluorescence, respectively. Data was analysed by Kruskal-Wallis test (qPCR delta CT values) and one-way analysis of variance (ANOVA) (immunofluorescence % area stained).

**Results:** Il-6 protein expression was significantly reduced in Tlr4 KO mice as compared to WT mice at 6 h in ileum and colon. mRNA expression was not significantly different between groups.

**Conclusion:** These findings support the hypothesis that Tlr4 deletion protects from irinotecan-induced mucositis by ameliorating Il-6 production. The effect appears to be post-transcriptionally mediated, although further research is required to determine if it is via a direct or indirect mechanism. In the future, Il-6 may be targeted therapeutically to ameliorate symptoms of mucositis.

## INTRODUCTION

Irinotecan is a DNA topoisomerase I-targeted chemotherapeutic agent, approved as a first line agent for metastatic colorectal cancer^1^. It causes exceptionally high rates of gastrointestinal (GI) mucositis in the form of severe diarrhoea, leading to delays in therapy, dose reductions or in severe cases, discontinuation of treatment^2^. The pivotal phase III clinical trials of irinotecan found it was associated with, severe diarrhoea in 20-35% of cases^3, 4^.

Irinotecan-related diarrhoea is biphasic. A cholinergic-type diarrhoea occurs within 24 hrs of administration and can be managed with atropine^5^. Of serious concern is the late-onset diarrhoea which occurs 24 h after irinotecan administration. It is associated with severe inflammation within the small and large intestine mucosa and lacks an effective management intervention. Although, there is limited information about the inflammatory events that lead to severe irinotecan-induced diarrhoea^6^; the interaction between the gut microbiome^7^ and Toll like receptor 4 (Tlr4) signalling^8^ has gained interest. Tlr4 is an innate immune receptor expressed on intestinal epithelial cells^9^ and immune cells of lamina propria, that exists in homeostasis with the intestinal environment^10^. Recent in vivo research has found Tlr4 to be a key mediator of irinotecan-induced mucositis^11^ and mice lacking Tlr4 are protected from severe irinotecan-induced mucositis^11^.

Irinotecan is metabolised to cytotoxic SN-38 by hepatic or GI carboxylesterase, which causes irreversible DNA damage. After glucuronidation by liver glucuronyltransferase (UGA1T1), inactive SN-38G is excreted into the intestine for elimination. However, in the intestinal lumen, bacterial β-glucuronidases regenerate toxic SN-38 from SN-38G^12^. This unique metabolic pathway not only results in high levels of intestinal toxicity, but also emphasises a key role played by gut microbiome^7^. Although irinotecan causes entire intestinal damage, ileum and colon are the most severely affected^13^. SN-38 is hypothesised to interact with Tlr4 at the MD-2 co-receptor binding pocket^14^. Tlr4/MD-2 receptor activation recruits adaptor protein MyD88 resulting in nuclear factor- κB (NF-κB) activation^15^. This causes downstream signalling and production of pro-inflammatory cytokines including tumour necrosis factor-α (Tnf-α), interleukin-1β (Il-1β), interleukin-6 (Il-6)^15^. Of this trio of cytokines, Il-6 is pivotal in setting of chronic inflammatory disease, including inflammatory bowel disease (IBD)^16^.

Il-6 is a pleiotropic cytokine that acts through the Il-6 receptor (Il-6R). Il-6R consists of two membrane bound non signalling α subunits and trans membranous signalling glycoprotein 130 (gp130). Binding of Il-6 to Il-6R recruits gp130 and initiates downstream signalling, including Janus kinases (JAKs) and phosphorylation of STAT3^17^. This pathway *(cis signalling)* promotes growth differentiation and regeneration. Only hepatocytes and some immune cells express Il-6Rα and thus respond to cis-signalling^16^. Metalloproteinases (ADAM17)^18^ cause proteolytic cleavage (shedding) of Il-6R from macrophages to create soluble form of Il-6R (sIl-6R). sIL-6R forms a complex with Il-6 (Il-6/sIl-6R) that can activate gp130, which is present on almost all body cells, hence, activating the cells which on their own are not responsive to Il-6 *(trans signalling)*^17^. This pathway is believed to mediate pro inflammatory characteristics of Il-6 via accumulation of T cells resistant to apoptosis and production of interferon-γ (IFN-γ), Tnf-α, and Il-1β^19^. As such, Tlr4-mediated upregulation of Il-6, and consequent activation of Il-6R may by a key driver of intestinal inflammation following exposure to irinotecan, although this has yet to be investigated directly.

## HYPOTHESES AND AIMS

In light of previous in vivo research, I hypothesise that “Tlr4 deletion protects against irinotecan-induced mucositis via regulation of Il-6 production”.

This hypothesis will be addressed by two distinct experimental aims

Aim 1: Quantify Tlr4 dependent Il-6 expression in irinotecan treated wild-type (WT) and Tlr4 knock out (KO) mice, at transcript and protein level in ileum and colon.

Aim 2: Quantify Il-6R expression at transcript and protein level in the ileum and colon of WT and Tlr4 KO mice.

## MATERIALS AND METHODS

### Ethics

This study utilised archival tissue from a previously conducted study approved by The University of Adelaide Animal Ethics Committee (M-2013-225). All experimental procedures compiled with the National Health and Medical Research Council (Australia) Code of Practice for Animal Care in Research and Training (2013).

### Experimental design

Female BALB/c-wild-type (WT) and BALB/c-*Tlr4*^-/-billy^ (Tlr4 KO) mice (n =48) weighing between 18 and 25 g (10–13 weeks) were used. The mice are described in detail else where^13^. Briefly, mice were given 270 mg/kg of intraperitoneal (i.p) irinotecan hydrochloride (provided by Pharmacia/Pfizer) in a sorbitol/lactic acid buffer (PH 3.4). Control mice were given the sorbitol/lactic acid buffer only. Atropine (0.03 mg/kg) was administered subcutaneously (s.c) to all mice prior to treatment to reduce the cholinergic diarrhoea. Mice were randomly assigned to treatment groups. Mice were anaesthetised using 200 mg/kg i.p. ilium sodium pentobarbital (60 mg/mL), and blood collected from the facial vein. Mice were killed at 6, 24, 48, 72 hours by transcardial perfusion with cold, sterile 1 × PBS (pH 7.4) followed by 4% paraformaldehyde (PFA) in 0.1 mol/L PBS (pH 7.4)^20^. Ileum and colon tissue of mice killed at 6 and 72 h were used in my project. Groups allocations are shown in Fig.1.

**Fig. 1.**
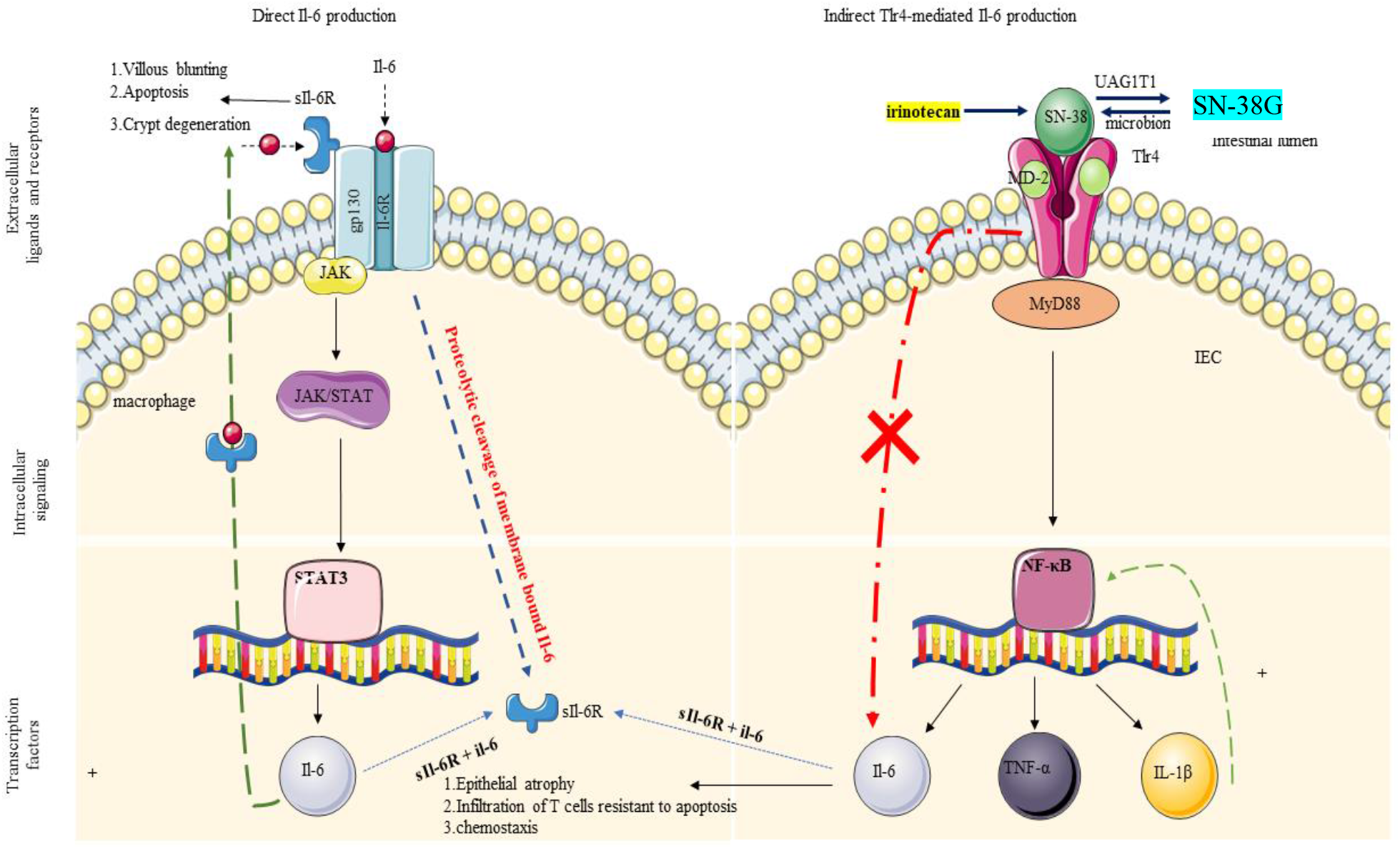
Pathways for IL-6 production. Direct IL-6 production occurs through the activation of a receptor formed by two distinct subunits, an alpha subunit for ligand specificity and glycoprotein (GP) 130. Binding of IL-6 to its receptor (IL-6R) initiates cellular events including JAK kinases, which when phosphorylated, activates signal transducers and activators of transcription-3 (STAT3). This then translocates to the nucleus resulting in the production of IL-6. Indirect IL-6 production is mediated through innate immune receptor, Toll-like receptor 4 (TLR4). TLR4 is expressed on glial, immune and intestinal epithelial cells with bacterial lipopolysaccharide (LPS) as its primary ligand. Upon LPS binding, TLR4 undergoes a conformational change resulting in the recruitment of TIR domains containing adaptor molecules. In particular, myeloid differentiation primary response (MyD)88-dependent signalling results in nuclear translocation of nuclear factor kappa B (NF-κB) and the subsequent production of IL-6^16^.

### RNA extraction

Total RNA was extracted from tissues using the NucleoSpin^®^ RNA II kit, as per manufacturer’s instructions (Macherey-Nagel, Germany). After series of filtration and washing steps, extracted RNA was eluted in sterile water and stored at −80°C. Total yield (ng/μl) and purity (260/280 ratio) were assessed by the BioTek Synergy Mx Microplate Reader (BioTek, USA), TAKE3 plate, and Gen5 (version 2.00.18) software.

### Reverse transcription and Real time PCR (qPCR)

cDNA was synthesised from 1μg of total RNA using the iScript cDNA Synthesis Kit (BioRad, Australia) as per manufacturer’s instructions. Briefly, RNA and RT master mix were incubated at varying temperatures in thermal cycler (Corbett research, Australia). Real time PCR was performed in total volume of 10 μL with 2 μL of cDNA (100 ng/μL), 4 μl of SYBR green fluorescence dye and 2 μl of each forward and reverse primers prediluted to 50 pmol/μl, using the Rotor-Gene 3000 (Corbett research, Australia). β-actin was used as the reference gene. Primers for Il-6 and Il-6R are listed in table 2.

**Table 3.**
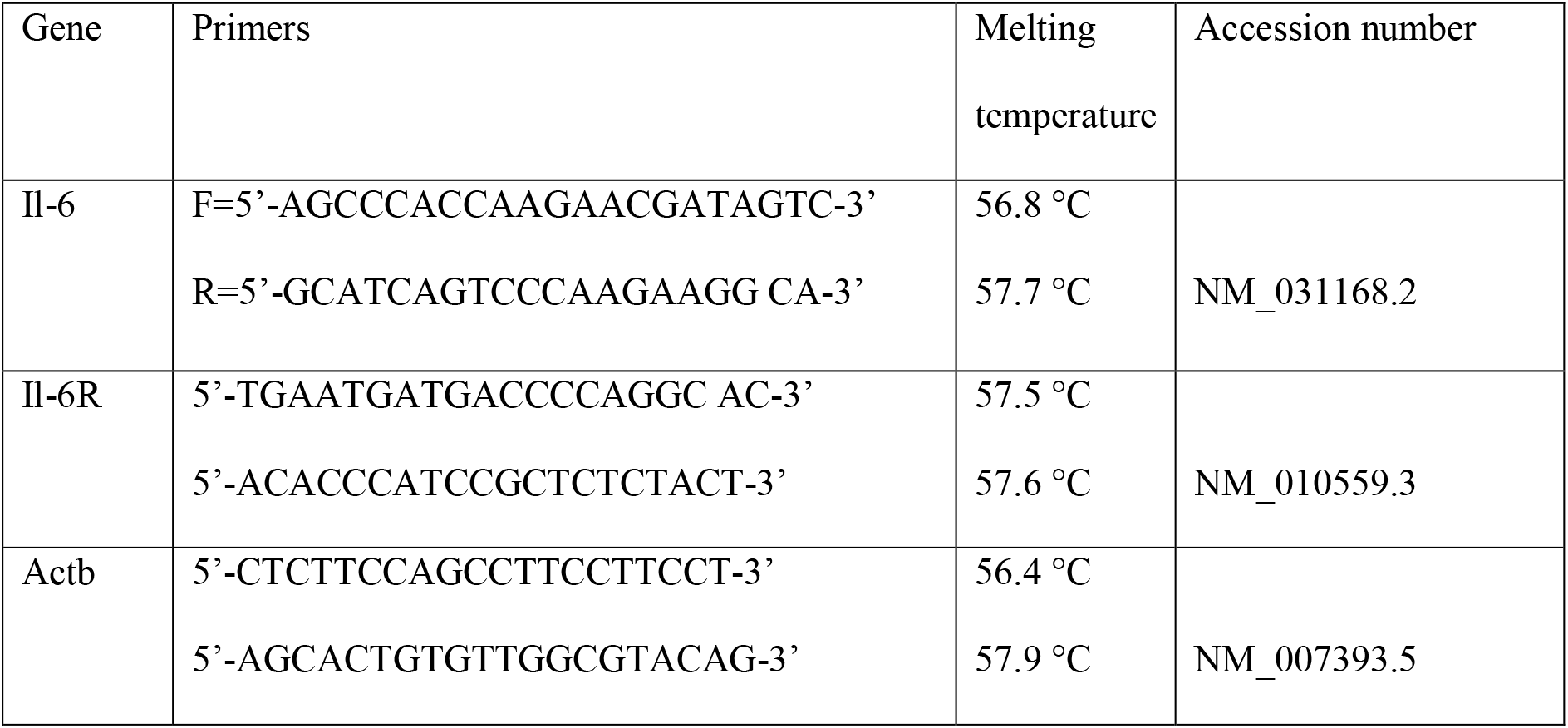
Primer sequence for Il-6, Il-6R

Two step PCR was run for Il-6 and Il-6R including activation at 95°C for 10 minutes, followed by 40 cycles of denaturation at 95°C for 15 seconds and annealing at 60°C for 45 seconds. All samples were run in triplicate with negative control (no cDNA template) added to each run. Mouse primers were designed using software Primer Blast-NCBI-NIH), efficacy checked through Net Primer (premier bio soft) and ordered from IDT (Targeted DNA technologies, Singapore). 2Δ*C*_T_^51^ method was used for relative quantification of both genes.

### Immunofluorescence (IF)

IF was used to identify protein localisation of Il-6 and Il-6R in formalin-fixed, paraffin-embedded tissue blocks ^11, 13^. Briefly, 5-μm sections of ileum, and colon were cut on microtome [Leica (RM 2235) Germany], and mounted onto FLEX IHC microscope slides (Dako, Denmark). Sections were deparaffinised in histolene and rehydrated through graded ethanol. Sections were immersed in EDTA-NaOH buffer (0.37 g/L EDTA, pH 9.0, preheated to 65°C) in Dako PT LINK (Dako; #PT101) and heated to 97°C for 20 minutes. Sections were incubated with 5% goat serum in PBS (Sigma-Aldrich Inc, St. Louis, MO) for 1 hour. The primary antibodies Il-6 [ Abcam (UK), ab 7737, 0.1 mg/ml, rabbit polyclonal anti-mouse antibody^21^, 1:100] and Il-6R [Bioss (UK), 1 μg/μl bs-1805R, rabbit polyclonal anti-mouse antibody^22^, 1:100] diluted in 5% NHS were applied for 1 hour. The specificity of Il-6^21^ and Il-6R^22^ antibodies has been assessed previously. Sections were then incubated with a fluorescently labelled secondary antibody [goat anti-rabbit secondary antibody, Alexa Fluor 488 conjugate, 2 mg/ml, 1:250, Invitrogen, USA] for 1 hour, diluted in 1× PBS + 1% BSA (Sigma-Aldrich, USA) and 2% FBS (Sigma-Aldrich, USA). Slides were washed in 1× PBS (PH 7.4), DAPI (4’,6-diamidino-2-phenylidole, 1 μg/mL, Sigma-Aldrich, Israel) was added followed by fluorshield (Sigma Aldrich, USA). Coverslip (Germany) was carefully mounted. Negative control (no primary antibody) was added to each run. Slides were visualised under the Olympus bx51 light microscope. IF was assessed for % staining^52^ in a blinded fashion by Image J (Fiji, version 2).

### Data analysis

Data were analysed using Prism version 8.0 (GraphPad Software, USA). IF data was log-transformed and normality confirmed by D’agostino-Pearson omnibus test. A one-way ANOVA with appropriate *post-hoc* testing was performed to identify statistical significance in normally distributed data. In other cases, a Kruskal–Wallis test with Dunn multiple comparisons test and Bonferroni correction were performed. *P* < 0.05 was considered significant.

## RESULTS

### Transcript expression

Relative quantification of Il-6 and Il-6R transcript expression in tissue homogenates was conducted by RT-PCR. No significant difference between treatment group or genotype was found for Il-6 at 6 or 72 h in ileum or colon (Fig 2).

**Figure 2.**
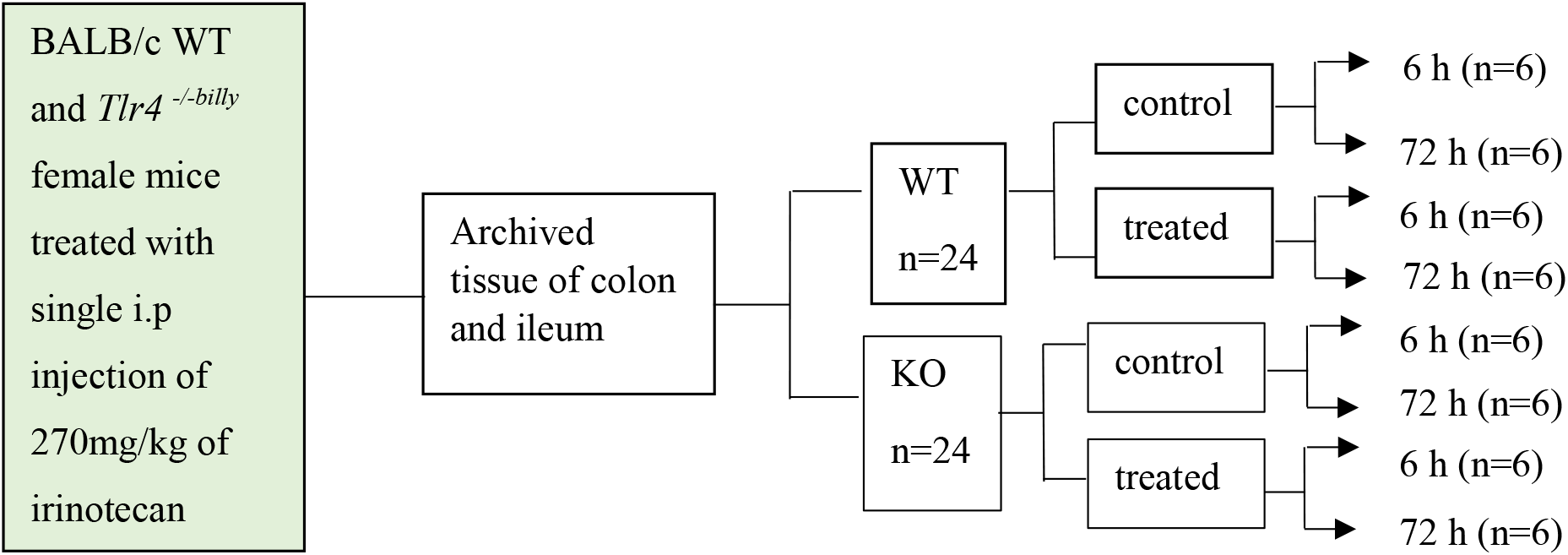
experimental plan

Similarly, no significant difference was found between treatment group or genotype for Il-6R at 6 or 72 h in ileum and colon (Fig.3).

### Protein expression

The tissue location and quantity of Il-6 and Il-6R protein was assessed by immunofluorescence. Positively stained cells were present in the lamina propria, blood vessels and peyer’s patches, while distinct epithelial staining was seen for Il-6R.

Within ileum tissue, irinotecan treatment caused a significant increase in Il-6 staining at 6 h in wild-type mice compared to controls. This effect was not observed in Tlr4 KO mice. Il-6 staining was also significantly less in irinotecan treated Tlr4 KO mice compared to wild type mice at 6 h There were no differences between groups at 72 h (Fig. 4).

**Fig. 4.**
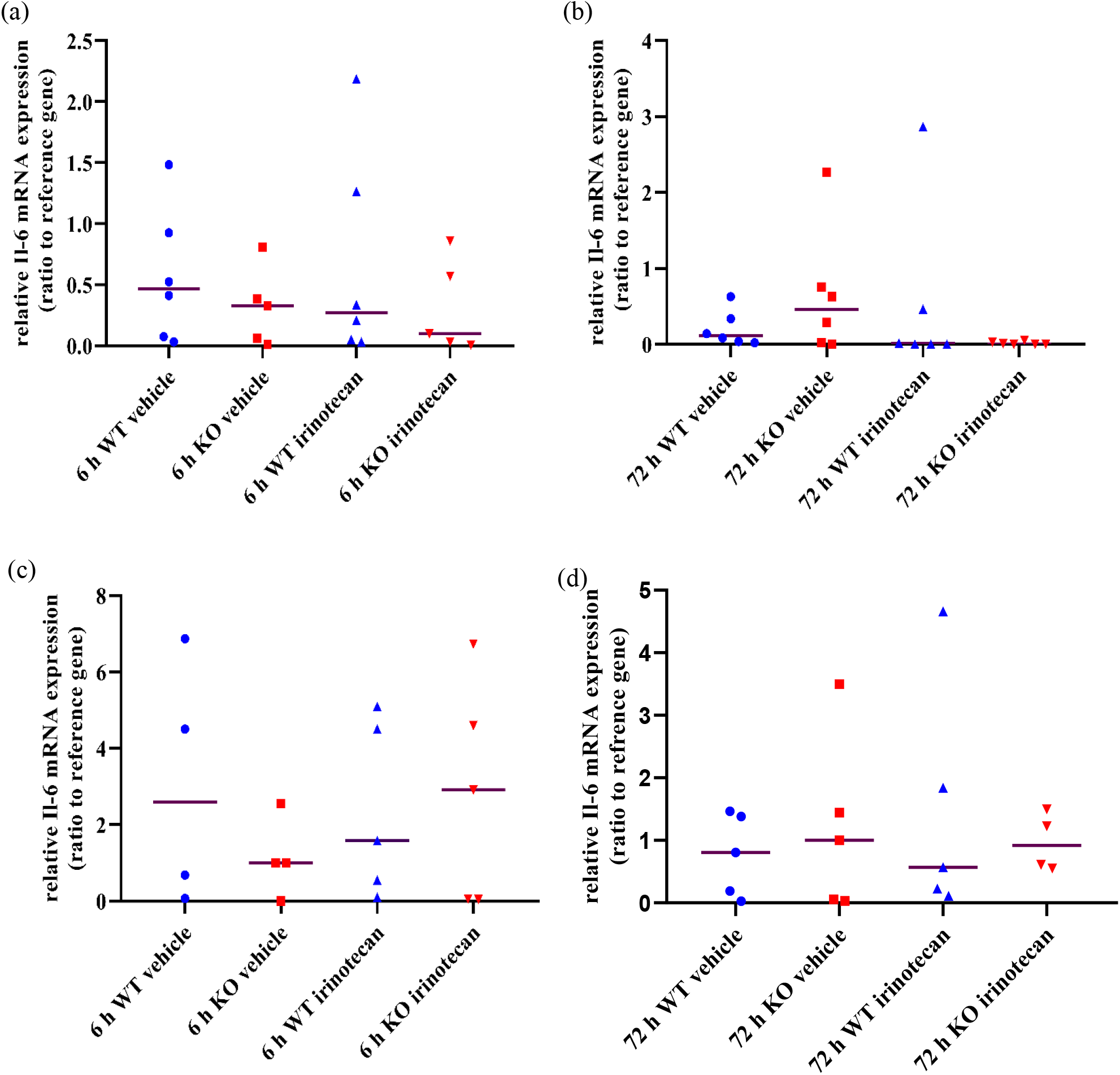
(a). Il-6 gene expression between WT and Tlr4 KO group in control and irinotecan-treated mice in ileum (a) 6 h. (b) 72 h and colon (c) 6 h. (d) 72 h. Data is shown as Il-6 expression relative to reference gene (β-actin), calculated by ΔCT. Line here is median. p = ns.

Within colon tissue, irinotecan treatment caused a significant increase in Il-6 staining at 6 h in wild-type mice compared to controls. This effect was not observed in Tlr4 KO mice. Il-6 staining was also significantly less in irinotecan treated Tlr4 KO mice compared to wild type mice at 6 h There were no differences between groups at 72 h (Fig.5)

**Fig. 5.**
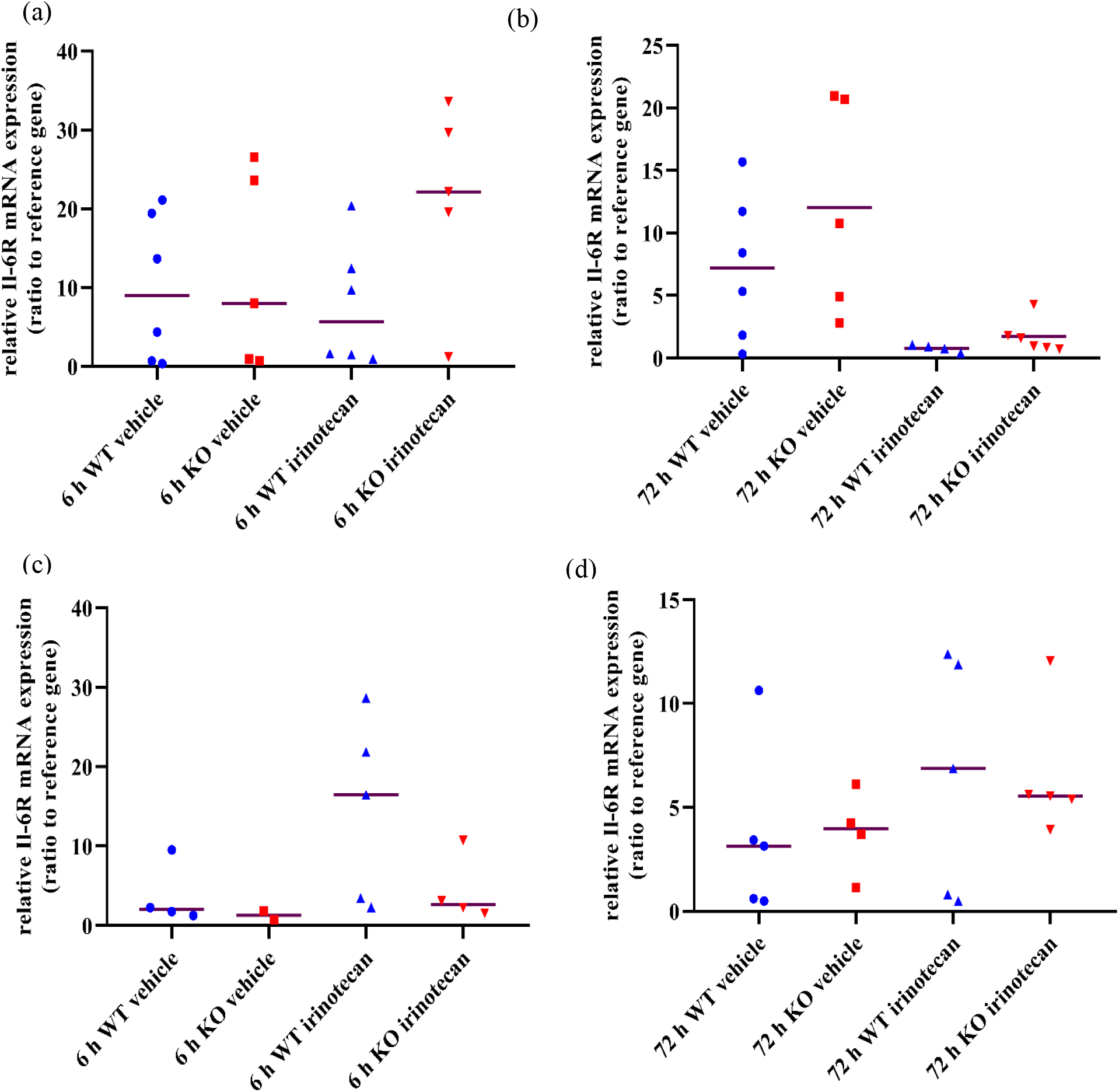
Il-6R gene expression between WT and Tlr4 KO group in control and irinotecan-treated mice in ileum (a) 6 h. (b) 72 h and colon (c) 6 h (d) 72 h. Data is shown as Il-6R expression relative to reference gene (β-actin), calculated by ΔCT. Line here is median. p = ns

No significant difference was seen for Il-6R protein expression between WT and Tlr4 KO groups at 6 h in ileum. While significant upregulation of Il-6R expression was seen in WT mice after treatment with irinotecan as compared to controls. (Fig.6)

**Fig. 6.**
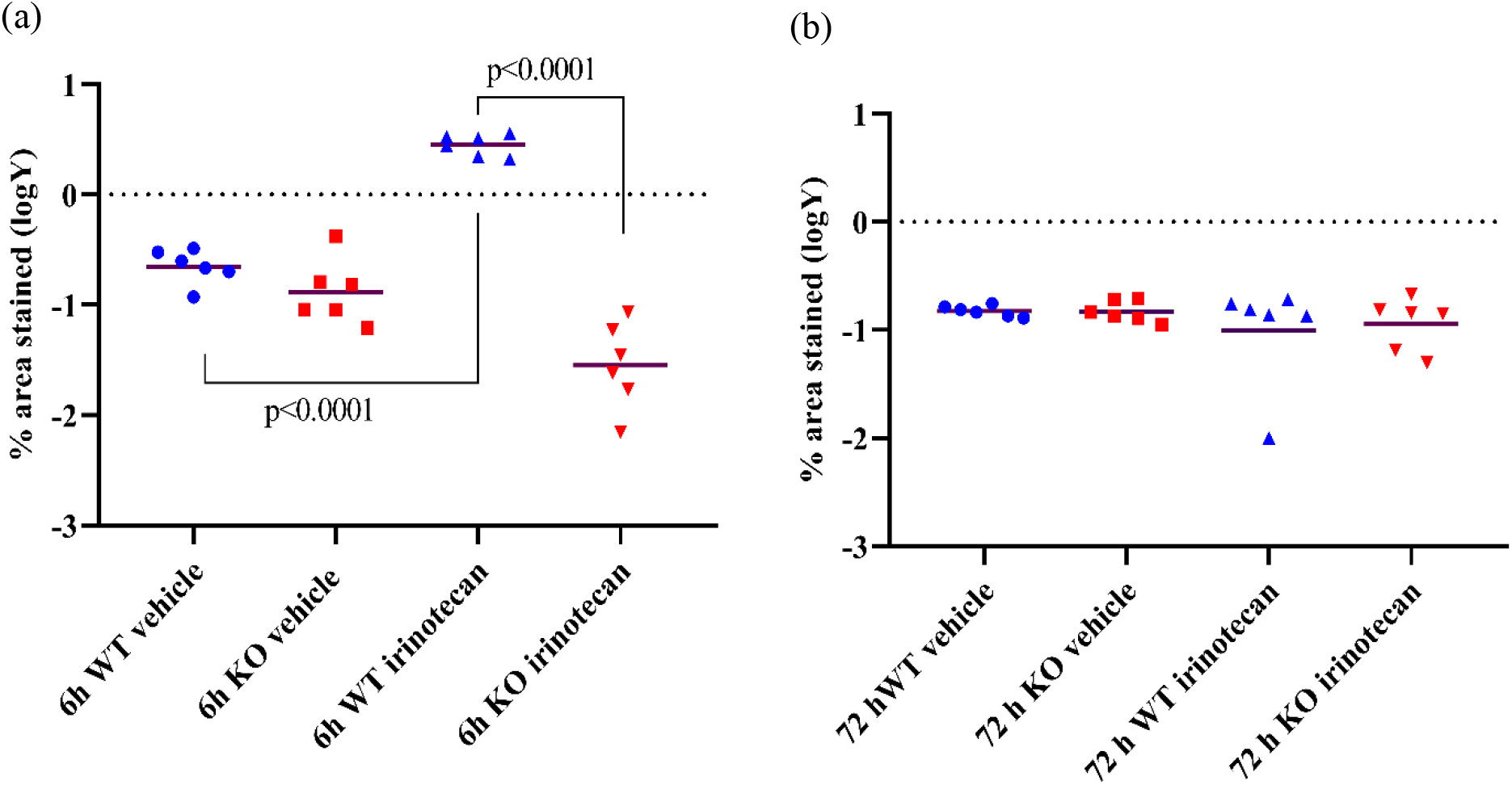
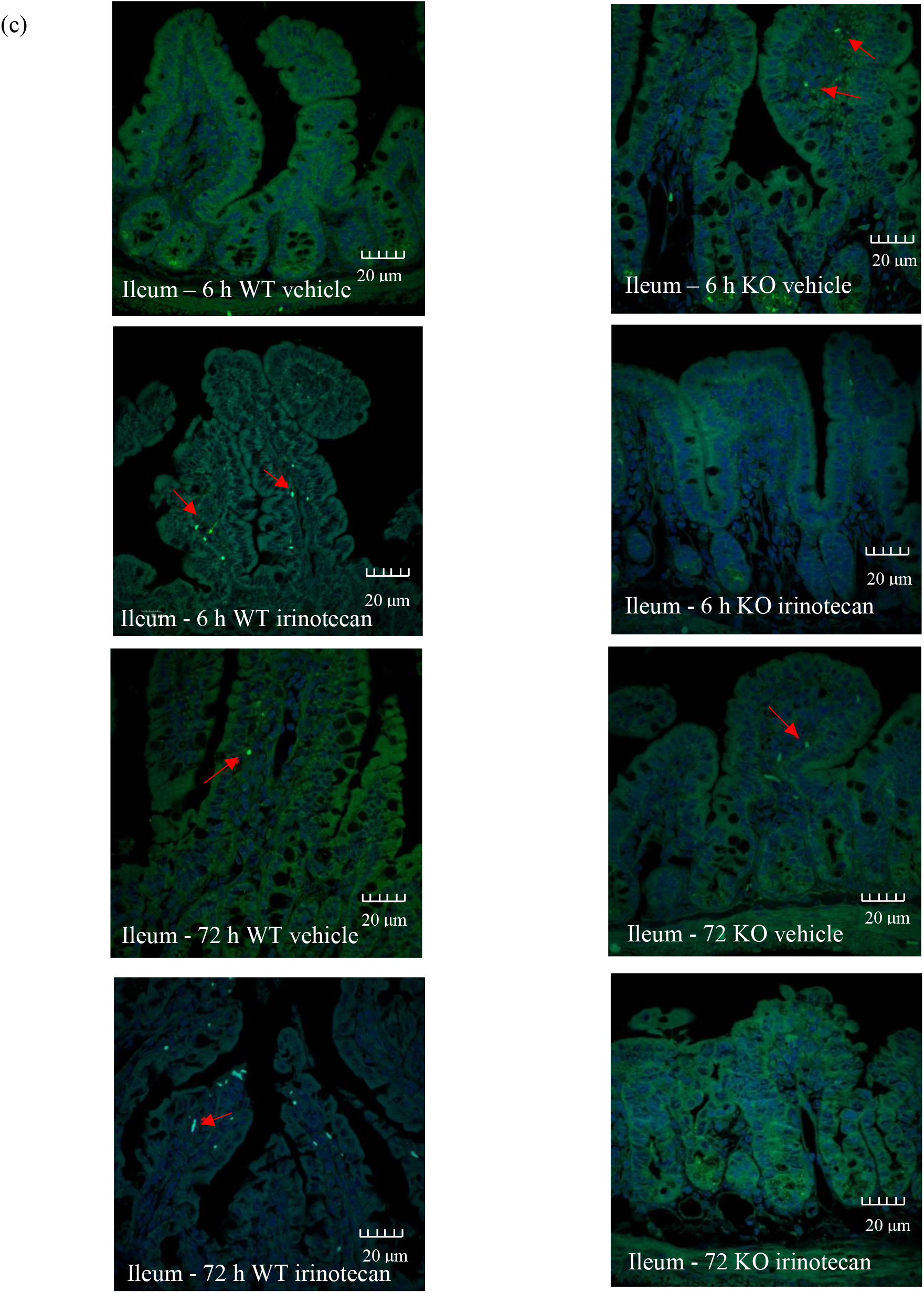
Il-6 protein expression between WT and Tlr4 KO group in control and treated mice in ileum. (a) at 6 h. (b) at 72 h. Data is shown as log transformation of percentage (%) stained area. Line here is mean. (c) Photomicrographs of representative Il-6 immunostaining at 40x. Arrow indicates positive staining.

Within colon tissue, irinotecan treatment caused a significant increase in Il-6R staining at 6 h in wild-type mice compared to controls. This effect was not observed in Tlr4 KO mice. Il-6R staining was also significantly less in irinotecan treated Tlr4 KO mice compared to wild type mice at 6 h There were no differences between groups at 72 h (Fig. 7)

**Fig. 7.**
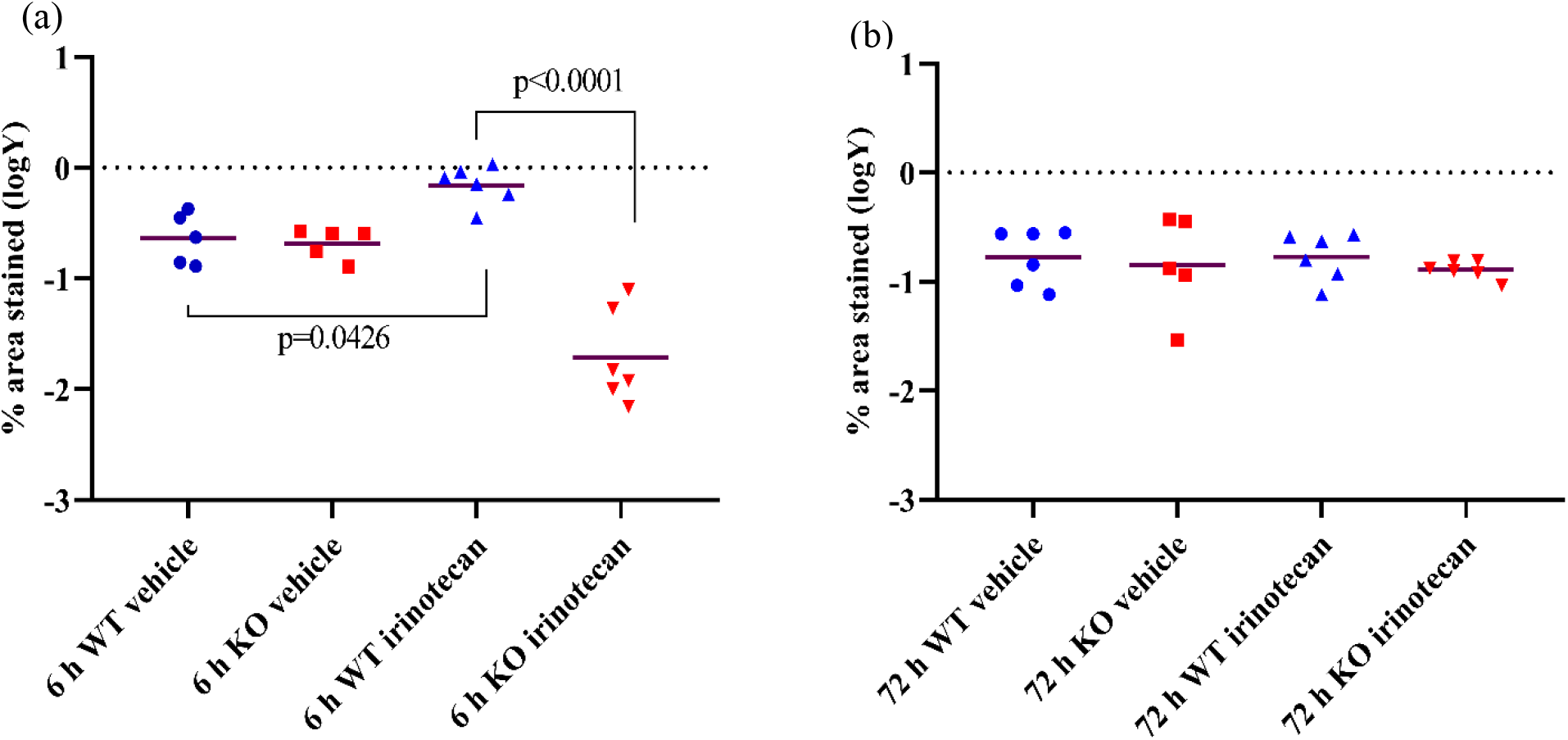
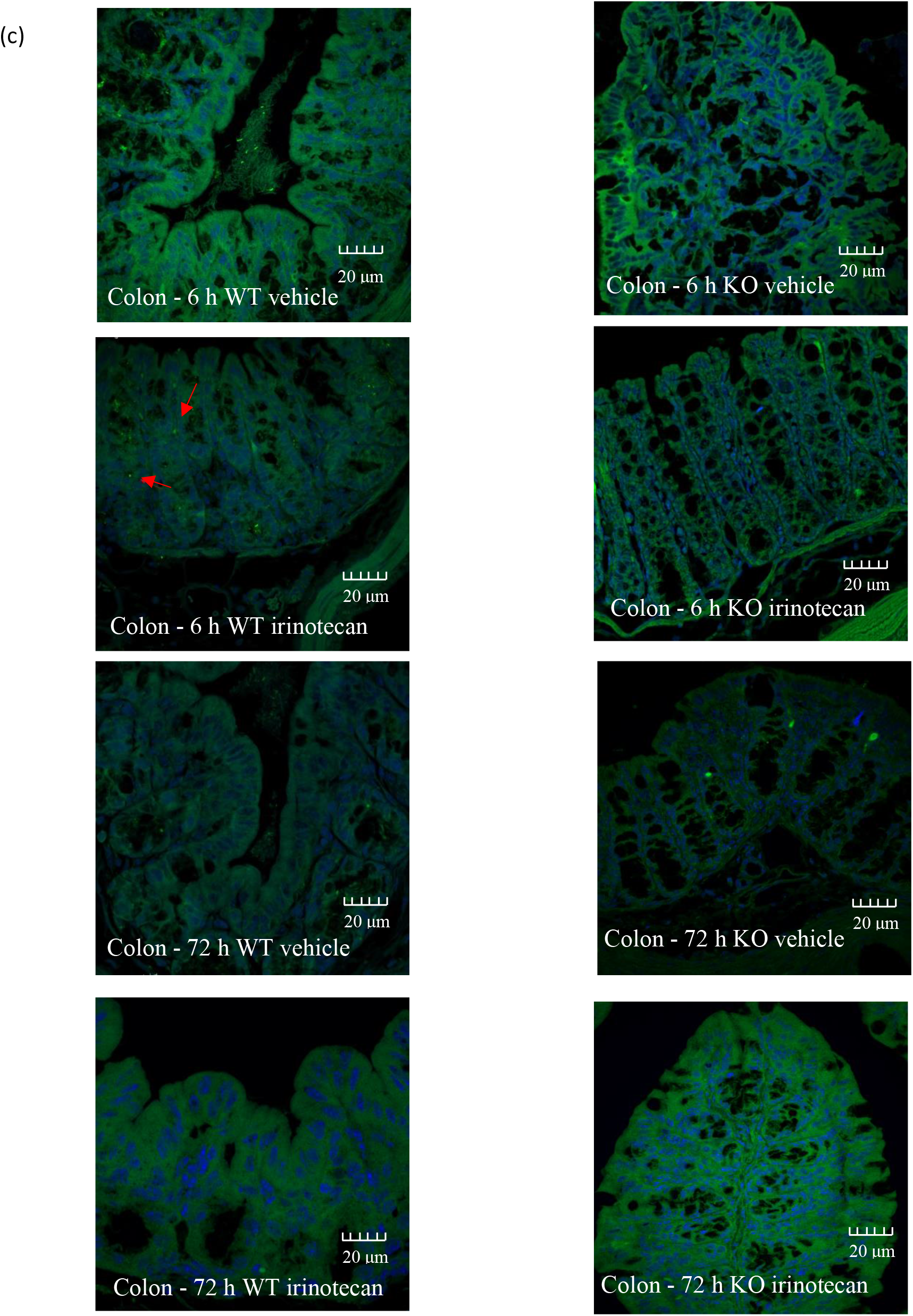
Il-6 protein expression between WT and Tlr4 KO group in control and treated mice in colon (a) 6 h. (b) 72 h. Data shown as log transformation of percentage (%) stained area. Line here is mean. (c) Photomicrographs of representative Il-6 immunostaining at 40x. Arrow indicate positive staining.

**Fig. 8.**
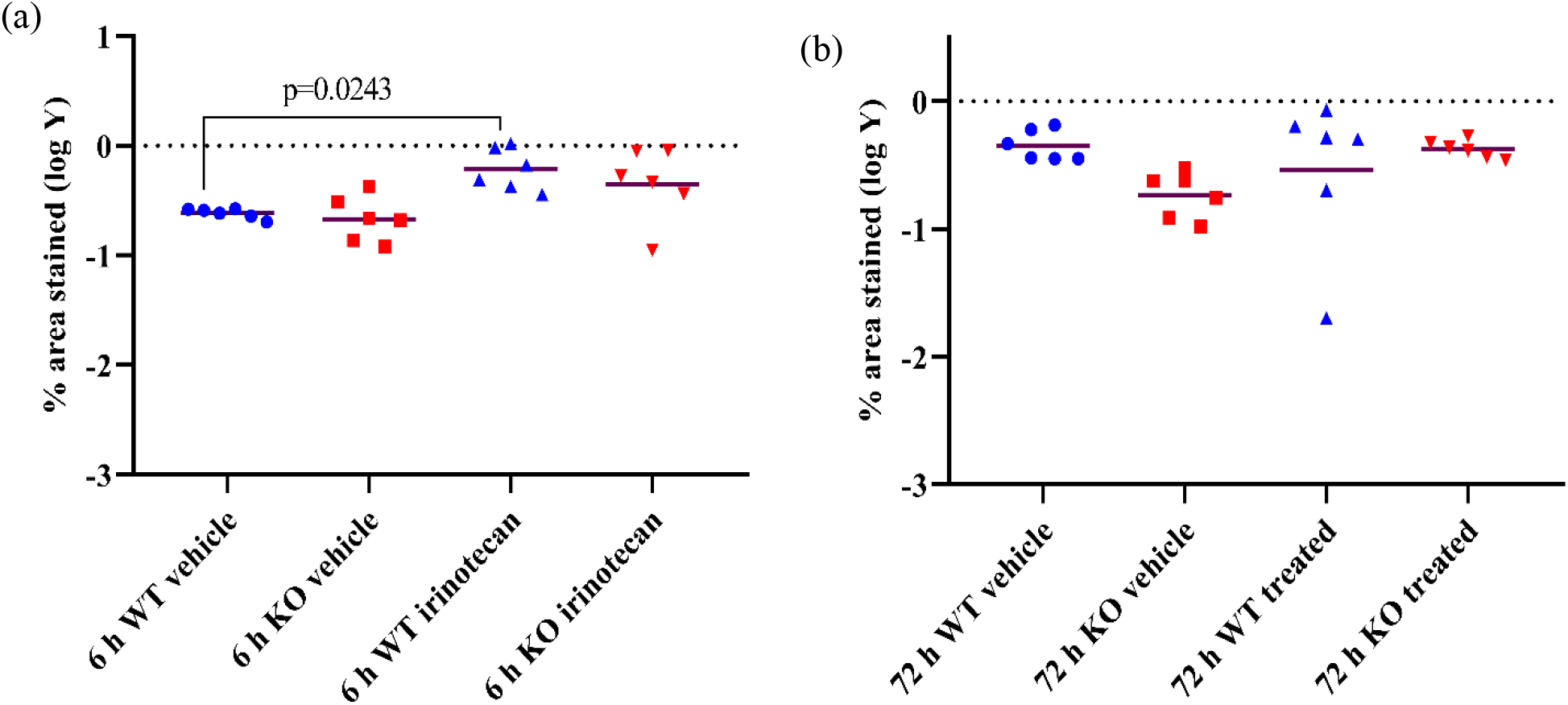
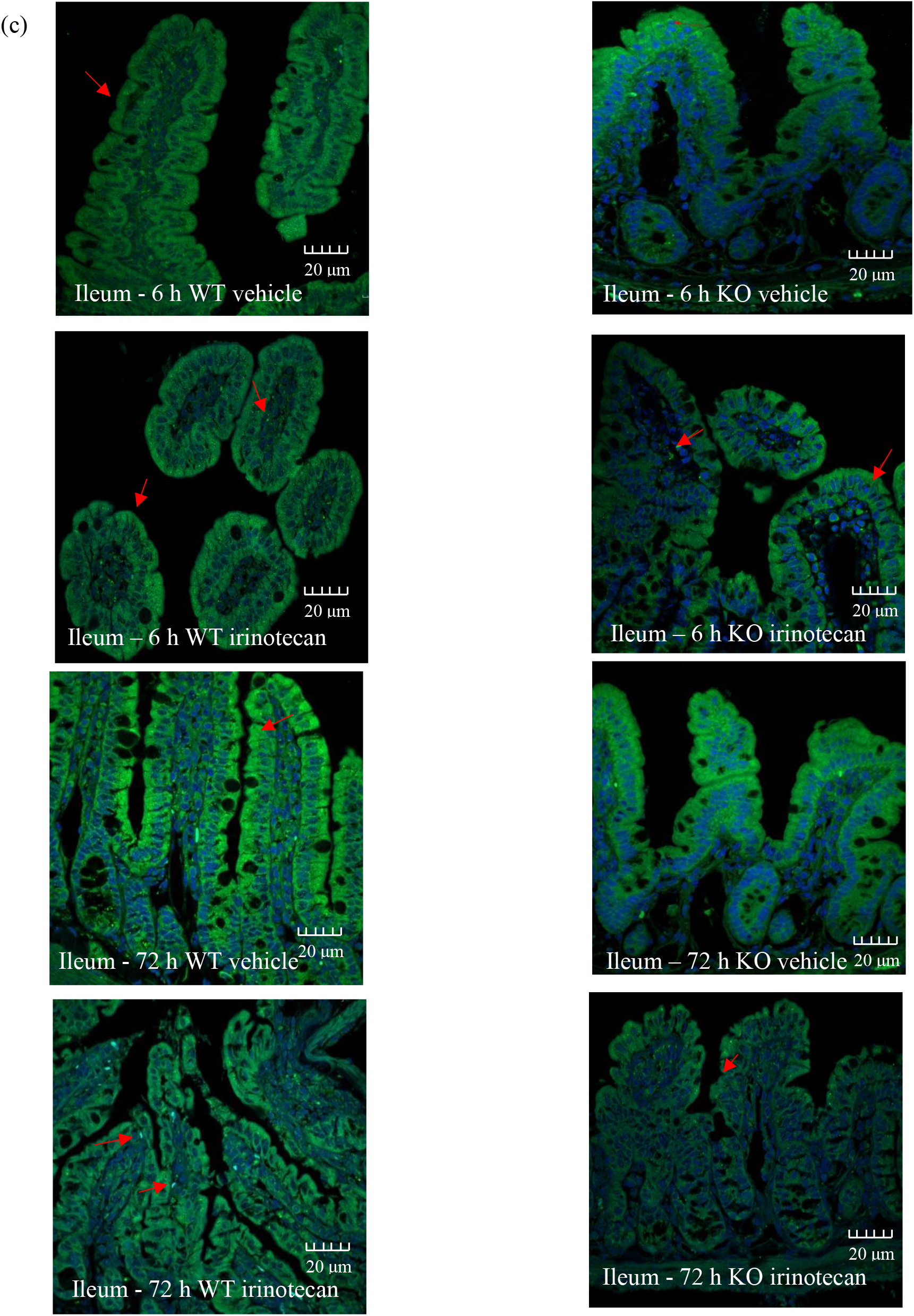
Il-6R protein expression between WT and Tlr4 KO group in control and treated mice in ileum. (a) at 6 h. (b) at 72 h. Data is shown as log transformation of percentage (%) stained area. Line here is mean. (c) Photomicrographs of representative Il-6R immunostaining at 40x. Arrow indicate positive staining.

**Fig. 9.**
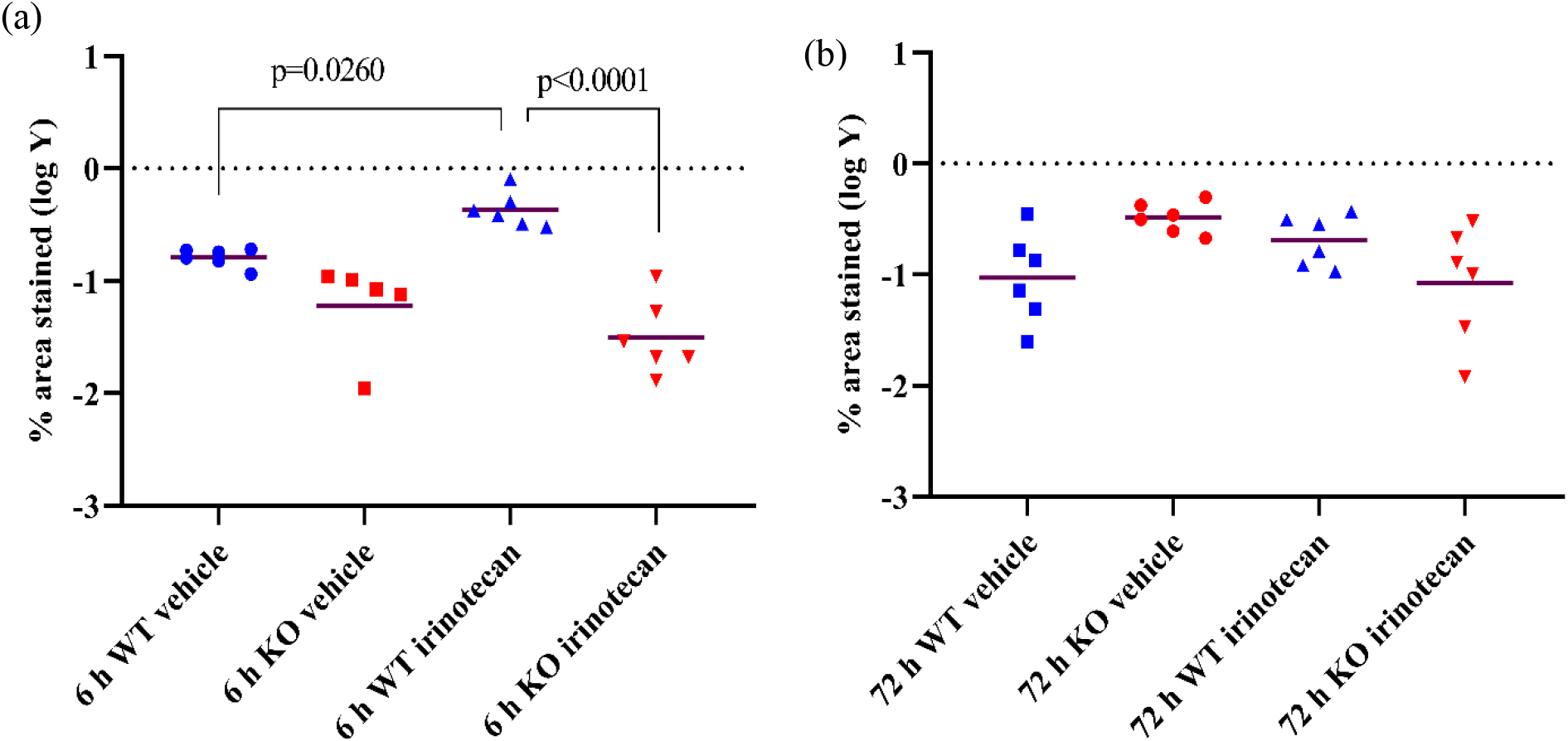
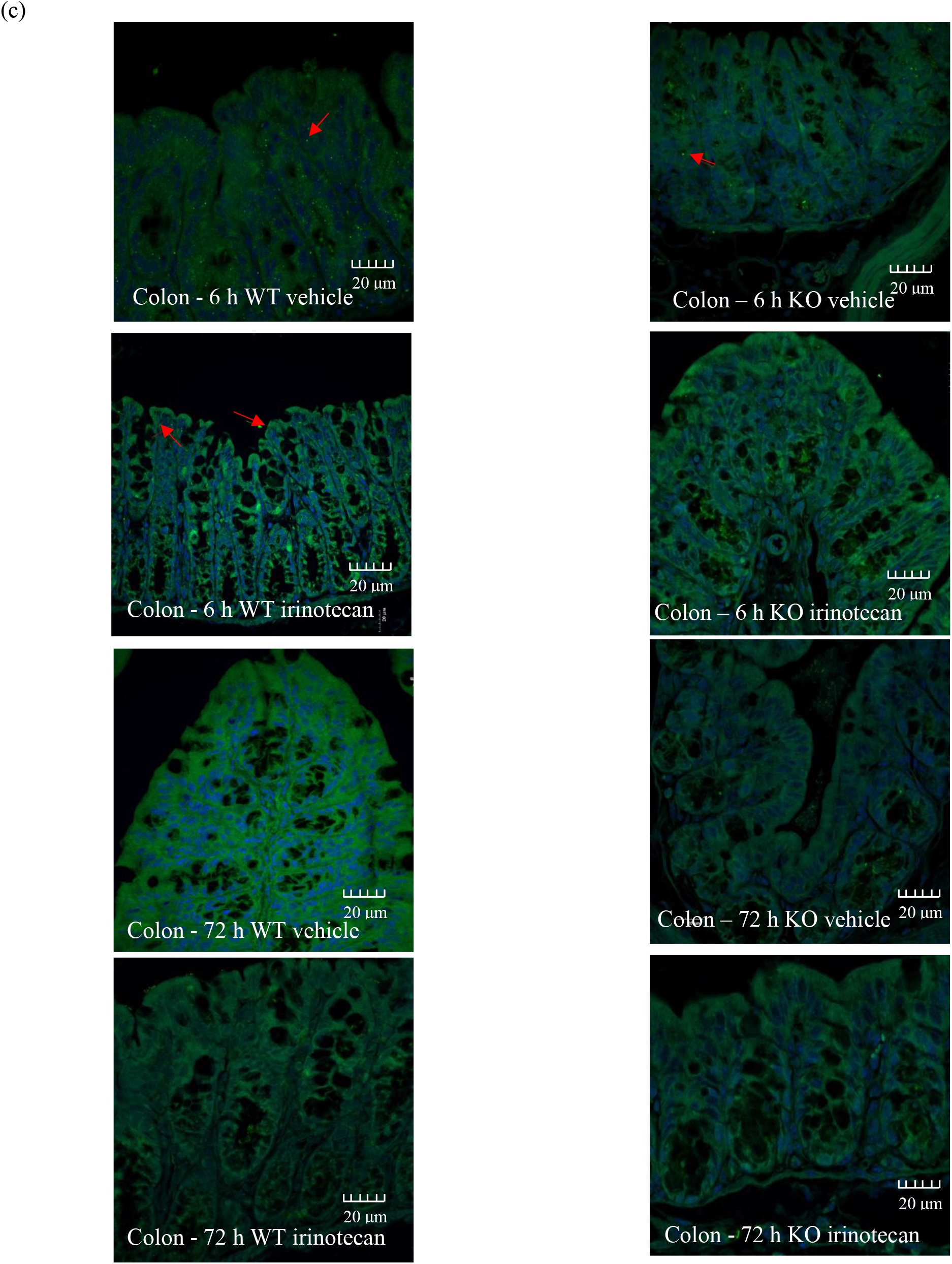
Il-6R protein expression between WT and Tlr4 KO group in control and treated mice in colon. (a).at 6 h. (b) at 72 h. Data is shown as log transformation of percentage (%) stained area. Line here is mean. (c) Photomicrographs of representative Il-6 immunostaining at 40x. arrow indicate positive staining.

## DISCUSSION

Mucositis in the form of severe diarrhoea is a debilitating adverse effect of irinotecan^2^. In previous studies, Tlr4 KO mice have shown reduced gut inflammation, Il-6 protein levels and diarrhoea compared to WT mice following irinotecan treatment^18^. As such, it was hypothesised that Il-6 up-regulation drives chemotherapy-induced mucositis and its production is mediated by Tlr4 activation. This study investigated the link between Tlr4 and Il-6 signalling in the context of irinotecan-induced mucositis.

Il-6 was significantly upregulated in WT irinotecan treated mice as compared to the control group at 6 h in ileum and colon. This finding is supported by previous research^13^, that showed significant Il-6 upregulation in WT irinotecan treated mice at 6 h^13^ and rats at 12 h^6^. While, Tlr4 KO mice in our study showed significantly reduced intestinal Il-6 protein expression compared to WT mice treated with irinotecan at 6 h as measured by immunofluorescence. Previous work had shown prevention of irinotecan-induced Il-6 upregulation using ELISA^13^. Although quantitative, ELISAs are limited in their interpretation due to the inability to visualise the location of protein expression. In contrast, immunofluorescene provides an opportunity to determine the location of Il-6 expression in the tissue structure. The cells expressing Il-6 as seen under confocal were located in lamina propria of ileum and colon, paeyers patches and blood vessels. This can be explained by the fact that Il-6 is secreted by macrophages produced from circulating monocytes, attracted to site of inflammation. While resident macrophages are responsible for maintaining intestinal homeostasis and rarely secrete pro inflammatory cytokines^23^.

Tlr4 activation has already been hypothesised to be a key driver of irinotecan induced mucositis that is targetable for clinical management^24^. Pre-clinical^25^ and clinical^26^ research indicated upregulated Tlr4 expression following irinotecan administration and radiotherapy respectively. Tlr4 KO mice had shown significant reductions in diarrhoea, weight loss^11^ and intestinal apoptosis as compared with WT mice after irinotecan administration^13^. On the contrary, a non-specific Tlr4 antagonist, (-)-naloxone, was not effective in reducing irinotecan-induced mucositis in Albino rats^24^. Moreover, Tlr4 antagonist, (-)-naloxone was associated with increased breast tumour growth and reduced irinotecan efficacy in Albino rats^24^. This emphasises the need for pharmacological inhibition of signalling products of Tlr4 for prevention of mucositis. Of the products of Tlr4 signalling, Il-6 has been found to be implicated in various inflammatory disorders especially IBD^27^.

In the view of current study, it can be hypothesised that interruption of Tlr4 signalling prevents signals that normally lead to Il-6 upregulation. Yet it is not clear if Tlr4 directly stimulates production of Il-6 from Il-6R-expressing macrophages and epithelial cells or indirectly by activation of NF-κB in cells responding to damage signals^19^. Although Tlr4 seems to regulate Il-6 production, an additional layer of complexity is expected to exist. The possibility arises because Tlr4 KO tissues in our study showed some degree of staining for Il-6, indicating that besides Tlr4, there may be some other factors responsible for Il-6 production. The five-phase model of mucositis presented by Sonis^15^ implies production of reactive oxygen species (ROS) as a first step in initiation of mucositis. Further research indicated that ROS is capable of activating NF-κB producing Il-6^28, 29^ and Tnf-α in alveolar macrophages^30^ and clinically inhibition of mitochondrial ROS production inhibits production of Il-6 and Tnf-α^31^. The impact of ROS on Il-6 production in mucositis has not been studied so far. This research provides solid foundation to determine if ROS constitutes an important stimulus in Il-6 production from macrophages in irinotecan induced mucositis. ROS also appears to exacerbate LPS mediated Il-6 production^28^; which is a potent Tlr4 agonist. Hence, a variety of different stimuli activate Il-6 which then serves as a chemo-attractant for other inflammatory cytokines. As such, while the results in this study and that of others support Tlr4-dependent Il-6 production, it is highly likely that Il-6 remains stimulated by other inflammatory pathways independent of Tlr4 activation.

While Il-6 production appears to be an important step in the initiation of mucositis, levels return to baseline by 72 h with no significant difference in the mRNA or protein expression of Il-6 in the ileum and colon between WT and KO mice at the later time point. This indicates that Il-6 is an initiator cytokine that is elevated in the pathway of irinotecan-induced mucositis, before other cytokines can enter the inflammatory cascade, highlighting a key point between direct cytotoxic damage and secondary indirect tissue injury. This is supported by preclinical^6, 32^ and clinical research^33^ that Il-6 appears in mucositis pathway as early as 6 h ^13^, causing infiltration of apoptosis resistant T cells and possibly serving as a chemo-attractant for Tnf-α and Il-1β producing cells that are elevated at 24 and 48 h respectively^13^. Il-1β and Tnf-α establish a positive feedback amplification of Il-6 production from NF-κB hence, potentiating the epithelial cell damage^16^. Blocking Il-6 production may also block production of other inflammatory cytokines by infiltrating immune cells. As such, therapeutically targeting Il-6 may an ideal approach to managing side effects of irinotecan without impacting on the direct cytotoxic properties of irinotecan and negating the challenges in specifically targeting intestinal Tlr4.

Il-6 inhibition has been investigated in other benign inflammatory disorders. The anti-Il-6 antibody, tocilizumab, has successfully been used in rheumatoid arthritis^34^. Moreover, in a clinical research, an Il-6 antibody (PF-04236921) has shown higher response and rates of remission in Crohn’s disease than placebo therapy^35^. Clinically, use of anti Il-6 monoclonal antibody has shown to reduce the severity of mucositis in multiple myeloma patients^36^. Yet further research is needed for potential use of anti Il-6 antibody in mucositis.

Talking about Il-6 signalling without mentioning Il-6R is not fair at all. So far, our researchers seem to be interested in sIl-6R owing to its pro-inflammatory nature^34^ and membrane bound Il-6R seems to be neglected for over a decade. So, in addition to tissue expression of Il-6, I also investigated the expression of its receptor Il-6R to explore its potential contribution to irinotecan induced mucositis. Il-6R is typically expressed on hepatocytes, spleen and immune cells^16^. Il-6 mediates its anti-inflammatory properties (regeneration and growth) by ***cis signalling***. while proinflammatory pathways are mediated via ***trans signalling***. As such, I also hypothesized that irinotecan induces mucositis is dependent on Tlr4 mediated Il-6R expression.

Irinotecan caused a significant increase in Il-6R expression in the ileum and colon. While the expression was significantly decreased in Tlr4 KO mice compared to WT controls in colon, I found no significant difference between two genotypes in ileum. The difference between regions may be explained by the underlying level of immune activation in colon related to increased microbial load^37^ and its macrophage population^38^. This can further be explained by the finding that colon has high Tlr4 expression as compared to ileum^39^. Importantly, I observed clear Il-6R expression on epithelial cells. This finding contradicts research that implies that Il-6R is expressed only on hepatocytes and macrophages^16^. However, research dating back two decades has indicated that this receptor was expressed on intestinal epithelial cells^40, 41^. Research in the late 90’s has indicated that Il-6R initially undetectable in foetal small bowel was expressed in intestine postnatally^42^. However, to the best of our knowledge there seems to be no latest follow up on this finding. More precisely, very few studies have shown a biological effect of Il-6 *cis signalling* on epithelial proliferation and mucosal barrier integrity in vitro, suggesting Il-6R is expressed on epithelial cells^43^. These findings therefore highlight a potential avenue for future research to delineate the expression and biological significance of epithelial Il-6R in intestinal homeostasis and inflammation. In addition to quantifiable expression, the localisation of Il-6R was affected by irinotecan. While untreated WT animals showed a small amount of epithelial Il-6R expression, owing to its role in crypt/villus maturation, growth and development^40^, Il-6R was preferentially expressed in the immune cells of lamina propria and blood stream.

In contrast, irinotecan-treated groups showed upregulation of epithelial Il-6R in ileum and colon. The epithelial expression which was distinct at 6 h in irinotecan treated mice seems to be regulated by Tlr4 in colon. Our finding is supported by previous research that indicates that LPS (a potent Tlr4 agonist) causes upregulation of membrane bound Il-6R on intestinal epithelium, while it downregulates the receptor on macrophages^40^, which seems to be result of shedding to produce sIl-6R for inflammatory reactions^44^. However, it is not clear why the epithelial Il-6R is upregulated in irinotecan treated mice. A possibility exists that, just like the paradoxical role of Tlr4 in acute^45^ and chronic inflammation^9^, epithelial Il-6R expression may be increased as a protective measure to combat mucositis. This is supported by evidence indicating the increase in intestinal perforations in rheumatic arthritis patients under therapy with tocilizumab, a human anti-Il-6R antibody, when compared to therapy with conventional disease-modifying anti -rheumatic drugs^46^. In a clinical phase II study testing the anti-Il-6 agent BMS-945429 in Crohn’s disease, two cases of intestinal perforations were reported resulting in premature termination of therapy (Clinicaltrials.gov Identifier: IM133-055). Similarly, in a murine model of colitis, administration of anti-Il-6 antibody enhanced intestinal damage^47^.

Together, this suggests that cis signalling may contribute to intestinal repair during inflammatory response; the regenerative and inflammatory processes may be going side by side in intestinal injury in which Il-6-dependent regeneration may be completely overridden by the inflammatory response. The presence of detectable Il-6 and Il-6R expression in non-inflamed intestinal tissues is of interest. This suggests that these proteins, in health, may be required to maintain homeostasis in the intestinal mucosa. Interestingly, besides Il-6R expressing on epithelium and in lamina propria cells, we found Il-6R expression in submucosal cells in ileum. Due to limited time and resources, we could not assess the type of cell, yet based on its shape and location, it is possibly the glia or ganglion of sub mucosal plexus. This possibility is supported by a study investigating link between Tlr4 and irinotecan induced pain^13^. Our study raises suspicion of Tlr4 mediated Il-6R expression, it is quite possible irinotecan induced pain might be regulated by Tlr4 mediated Il-6R expression in enteric nervous system. Another study indicates the expression of Il-6 in enteric glia cells in rats^48^. Moreover, inhibition of Il-6R in vivo in the irritable bowel syndrome rat model reduced visceral pain sensitivity^49^ indicating the Il-6R expression on in enteric glia cells. In order to understand this mechanism in more detail and identify translatable aspects for pain management, further preclinical research is required.

Besides the protein expression, Il-6 and Il-6R mRNA showed no significant difference between genotypes or treatments at either the early or later time point in the ileum and colon. This finding contradicts with our results for protein expression. The possible reason may be that changes in Il-6 and Il-6R transcript occurred before the tissue was collected. This can be further explained by fact that different genes are upregulated at different time points in response to stress, expressing as early as 1 h^50^. Other mechanisms like post-translation processing, reduced internalisation and breakdown of Il-6R, changes in the ratio of soluble to membrane-bound Il-6R and gp130 levels might also be controlling the measurable increase in Il-6 and Il-6R. However, future study needs to be done to explore these potential mechanisms.

Importantly, the concept of Tlr4 mediated Il-6 production in irinotecan induced mucositis remains speculative and therefore warrants further research to determine dual role of mysterious Il-6R in intestinal mucositis and its regulation by Tlr4. Similarly, the conclusions drawn in this study must be interpreted with appropriate recognition of the study limitations. For example, our study used archived samples and RT PCR is on whole tissue, while using mucosal scraping may have been more informative to determine epithelial expression of Il-6R. Moreover, our study provides no information about shedding of Il-6R to sIl-6R. Future research elucidating the cells responsible for shedding of Il-6R and production of Il-6 could be achieved by flow cytometry. We can extend our research in future by using colorectal tumour bearing mice to investigate effects of Tlr4 deletion on irinotecan induced mucositis. Previously, clinical research has indicated that Il-6 is accompanied by a higher expression in intestinal epithelium of CRC patients as compared to controls^41^.

## CONCLUSION

Our results support previous findings that Tlr4 KO mice have an impaired Il-6 response and reduced diarrhoea suggesting that targeting Il-6 may be more sensible approach to prevent irinotecan-induced mucositis. As such, we suggest that IL-6 may be a more promising therapeutic target to prevent or reduce the severity of GI mucositis caused by irinotecan and should form the basis of new research models.

## Acknowledgements

I would like to thank Associate Professor Joanne M Bowen and Dr Hannah Wardill for their ongoing support and dedication for this research. This project is funded by Ray and Shirl Norman cancer research trust.

## REFERENCES

1. Fujita K, Kubota Y, Ishida H & Sasaki Y (2015). Irinotecan, a key chemotherapeutic drug for metastatic colorectal cancer. World J Gastroentero. 21, 12234–12248.

2. Lalla RV & Peterson DE (2006). Treatment of mucositis, including new medications. Cancer J. 12, 348–354.

3. Saltz LB, Cox JV, Blanke C, Rosen LS, Fehrenbacher L, Moore MJ, Maroun JA, Ackland SP, Locker PK & Pirotta N (2000). Irinotecan plus fluorouracil and leucovorin for metastatic colorectal cancer. New Engl J Med. 343, 905–914.

4. Michael M, Brittain M, Nagai J, Feld R, Hedley D, Oza A, Siu L & Moore MJ (2004). Phase II study of activated charcoal to prevent irinotecan-induced diarrhea. J Clin Oncol. 22, 4410–4417.

5. Logan RM, Stringer AM, Bowen JM, Yeoh AS, Gibson RJ, Sonis ST & Keefe DM (2007). The role of pro-inflammatory cytokines in cancer treatment-induced alimentary tract mucositis: pathobiology, animal models and cytotoxic drugs. Cancer Treat Rev. 33, 448–460.

6. Logan RM, Gibson RJ, Bowen JM, Stringer AM, Sonis ST & Keefe DM (2008). Characterisation of mucosal changes in the alimentary tract following administration of irinotecan: implications for the pathobiology of mucositis. Cancer Chemother Pharmacol. 62, 33–41.

7. Stringer AM, Gibson RJ, Logan RM, Bowen JM, Yeoh AS & Keefe DM (2008). Faecal microflora and beta-glucuronidase expression are altered in an irinotecan-induced diarrhea model in rats. Cancer biol ther. 7, 1919–1925.

8. Wardill HR, Gibson RJ, Logan RM & Bowen JM (2014). TLR4/PKC-mediated tight junction modulation: A clinical marker of chemotherapy-induced gut toxicity? Inter J Cancer. 135, 2483–2492.

9. Cario E, Rosenberg IM, Brandwein SL, Beck PL, Reinecker HC & Podolsky DK (2000). Lipopolysaccharide activates distinct signaling pathways in intestinal epithelial cell lines expressing Toll-like receptors. J immunol. 164, 966–972.

10. Chabot S, Wagner JS, Farrant S & Neutra MR (2006). TLRs regulate the gatekeeping functions of the intestinal follicle-associated epithelium. J immunol. 176, 4275–4283.

11. Wardill H, Secombe K, Van Sebille Y, White I, Gibson R, Logan R & Bowen J (2015). TLR4 deletion attenuates irinotecan-induced gut toxicity and barrier dysfunction in the BALB/CMOUSE offering a new therapeutic target. Support Care Cancer. 23, S105–S106.

12. Tallman MN, Miles KK, Kessler FK, Nielsen JN, Tian X, Ritter JK & Smith PC (2007). The contribution of intestinal UDP-glucuronosyltransferases in modulating 7-ethyl-10-hydroxy-camptothecin (SN-38)-induced gastrointestinal toxicity in rats. J Pharmacol Exp Ther. 320, 29–37.

13. Wardill HR, Gibson RJ, Van Sebille YZA, Secombe KR, Coller JK, White IA, Manavis J, Hutchinson MR, Staikopoulos V, Logan RM & Bowen JM (2016). Irinotecan-Induced gastrointestinal dysfunction and pain are mediated by common TLR4-dependent mechanisms. Mol Cancer Ther. 15, 1376–1386.

14. Wong DVT, Ribeiro-Filho HV, Wanderley CWS, Leite C, Lima JB, Assef ANB, Cajado AG, Batista GLP, Gonzalez RH, Silva KO, Borges LPC, Alencar NMN, Wilke DV, Cunha TM, Figueira ACM, Cunha FQ & Lima-Junior RCP (2019). SN-38, the active metabolite of irinotecan, inhibits the acute inflammatory response by targeting toll-like receptor 4. Cancer Chemother Pharmacol. 83, 1–12.

15. Sonis ST (2004). Oral mucositis in cancer therapy. J Support Oncol. 2, 3–8.

16. Naugler WE & Karin M (2008). The wolf in sheep’s clothing: the role of interleukin-6 in immunity, inflammation and cancer. Trends Mol Med. 14, 109–119.

17. Rose-John S (2012). IL-6 trans-signaling via the soluble IL-6 receptor: importance for the pro-inflammatory activities of IL-6. Int J Biol Sci. 8, 1237–1247.

18. Arnold P, Boll I, Rothaug M, Schumacher N, Schmidt F, Wichert R, Schneppenheim J, Lokau J, Pickhinke U, Koudelka T, Tholey A, Rabe B, Scheller J, Lucius R, Garbers C, Rose-John S & Becker-Pauly C (2017). Meprin Metalloproteases Generate Biologically Active Soluble Interleukin-6 Receptor to Induce Trans-Signaling. Sci Rep. 7, 44053.

19. Karin M, Lawrence T & Nizet V (2006). Innate immunity gone awry: linking microbial infections to chronic inflammation and cancer. Cell. 124, 823–835.

20. Wardill HR, Gibson RJ, Van Sebille YZ, Secombe KR, Coller JK, White IA, Manavis J, Hutchinson MR, Staikopoulos V, Logan RM & Bowen JM (2016). Irinotecan-Induced Gastrointestinal Dysfunction and Pain Are Mediated by Common TLR4-Dependent Mechanisms. Mol Cancer Ther. 15, 1376–1386.

21. Park JS, Choi J, Kwon JY, Jung KA, Yang CW, Park SH & Cho ML (2018). A probiotic complex, rosavin, zinc, and prebiotics ameliorate intestinal inflammation in an acute colitis mouse model. J Transl Med. 16, 37.

22. Urushima H, Fujimoto M, Mishima T, Ohkawara T, Honda H, Lee H, Kawahata H, Serada S & Naka T (2017). Leucine-rich alpha 2 glycoprotein promotes Th17 differentiation and collagen-induced arthritis in mice through enhancement of TGF-beta-Smad2 signaling in naive helper T cells. Arthritis Res Ther. 19, 137.

23. Hashimoto D, Chow A, Noizat C, Teo P, Beasley MB, Leboeuf M, Becker CD, See P, Price J & Lucas D (2013). Tissue-resident macrophages self-maintain locally throughout adult life with minimal contribution from circulating monocytes. Immunity. 38, 792–804.

24. Coller JK, Bowen JM, Ball IA, Wardill HR, van Sebille YZ, Stansborough RL, Lightwala Z, Wignall A, Shirren J, Secombe K & Gibson RJ (2017). Potential safety concerns of TLR4 antagonism with irinotecan: a preclinical observational report. Cancer Chemother Pharmacol. 79, 431–434.

25. Bowen J, Coller J, Hutchinson M & Gibson R (2012). Expression of TLRs in the rat intestine following chemotherapy for cancer. Brain Behav Immun. 26, S27.

26. Guney Y, Ozel Turkcu U, Hicsonmez A, Nalca Andrieu M & Kurtman C (2007). Ghrelin may reduce radiation-induced mucositis and anorexia in head-neck cancer. Med Hypotheses. 68, 538–540.

27. Yamamoto M, Yoshizaki K, Kishimoto T & Ito H (2000). IL-6 is required for the development of Th1 cell-mediated murine colitis. J Immunol. 164, 4878–4882.

28. Naik E & Dixit VM (2011). Mitochondrial reactive oxygen species drive proinflammatory cytokine production. J Exp Med. 208, 417–420.

29. Takada Y, Mukhopadhyay A, Kundu GC, Mahabeleshwar GH, Singh S & Aggarwal BB (2003). Hydrogen peroxide activates NF-kappa B through tyrosine phosphorylation of I kappa B alpha and serine phosphorylation of p65: evidence for the involvement of I kappa B alpha kinase and Syk protein-tyrosine kinase. J Biol Chem. 278, 24233–24241.

30. Simeonova PP & Luster MI (1995). Iron and reactive oxygen species in the asbestos-induced tumor necrosis factor-alpha response from alveolar macrophages. Am J Resp cell Mol. 12, 676–683.

31. Bulua AC, Simon A, Maddipati R, Pelletier M, Park H, Kim KY, Sack MN, Kastner DL & Siegel RM (2011). Mitochondrial reactive oxygen species promote production of proinflammatory cytokines and are elevated in TNFR1-associated periodic syndrome (TRAPS). J Exp Med. 208, 519–533.

32. Logan RM, Stringer AM, Bowen JM, Gibson RJ, Sonis ST & Keefe DM (2008). Serum levels of NFkappaB and pro-inflammatory cytokines following administration of mucotoxic drugs. Cancer Biol Ther. 7, 1139–1145.

33. Logan RM, Gibson RJ, Sonis ST & Keefe DM (2007). Nuclear factor-kappaB (NF-kappaB) and cyclooxygenase-2 (COX-2) expression in the oral mucosa following cancer chemotherapy. Oral Oncol. 43, 395–401.

34. Diaz-Torne C, Ortiz MDA, Moya P, Hernandez MV, Reina D, Castellvi I, De Agustin JJ, Fuente D, Corominas H, Sanmarti R, Zamora C, Canto E & Vidal S (2018). The combination of IL-6 and its soluble receptor is associated with the response of rheumatoid arthritis patients to tocilizumab. Semin Arthritis Rheu. 47, 757–764.

35. Danese S, Vermeire S, Hellstern P, Panaccione R, Rogler G, Fraser G, Kohn A, Desreumaux P, Leong R & Comer G (2016). Results of ANDANTE, a randomised clinical study with an anti-IL6 antibody (PF-04236921) in subjects with Crohn’s disease who are antitumour necrosis factor inadequate responders. J Crohns Colitis. 10, S12–S13.

36. Rossi JF, Fegueux N, Lu ZY, Legouffe E, Exbrayat C, Bozonnat MC, Navarro R, Lopez E, Quittet P, Daures JP, Rouille V, Kanouni T, Widjenes J & Klein B (2005). Optimizing the use of anti-interleukin-6 monoclonal antibody with dexamethasone and 140 mg/m2 of melphalan in multiple myeloma: results of a pilot study including biological aspects. Bone Marrow Transpl. 36, 771–779.

37. Ahmed S, Macfarlane GT, Fite A, McBain AJ, Gilbert P & Macfarlane S (2007). Mucosa-associated bacterial diversity in relation to human terminal ileum and colonic biopsy samples. Appl Environ Microbiol 73, 7435–7442.

38. Mahida Y, Patel S, Gionchetti P, Vaux D & Jewell D (1989). Macrophage subpopulations in lamina propria of normal and inflamed colon and terminal ileum. Gut. 30, 826–834.

39. Price AE, Shamardani K, Lugo KA, Deguine J, Roberts AW, Lee BL & Barton GM (2018). A map of Toll-like receptor expression in the intestinal epithelium reveals distinct spatial, cell type-specific, and temporal patterns. Immunity. 49, 560–575.

40. Panja A, Goldberg S, Eckmann L, Krishen P & Mayer L (1998). The regulation and functional consequence of proinflammatory cytokine binding on human intestinal epithelial cells. J Immunol. 161, 3675–3684.

41. Shirota K, LeDuy L, Yuan SY & Jothy S (1990). Interleukin-6 and its receptor are expressed in human intestinal epithelial cells. Virchows Arch B. 58, 303–308.

42. Dame JB & Juul SE (2000). The distribution of receptors for the pro-inflammatory cytokines interleukin (IL)-6 and IL-8 in the developing human fetus. Early Hum Dev. 58, 25–39.

43. Wang L, Walia B, Evans J, Gewirtz AT, Merlin D & Sitaraman SV (2003). IL-6 induces NF-kappa B activation in the intestinal epithelia. J Immunol. 171, 3194–3201.

44. Briso EM, Dienz O & Rincon M (2008). Soluble IL-6R is produced by IL-6R ectodomain shedding in activated CD4 T cells. Journal of immunology. 180, 7102–7106.

45. Fukata M, Michelsen KS, Eri R, Thomas LS, Hu B, Lukasek K, Nast CC, Lechago J, Xu R, Naiki Y, Soliman A, Arditi M & Abreu MT (2005). Toll-like receptor-4 is required for intestinal response to epithelial injury and limiting bacterial translocation in a murine model of acute colitis. Am J Physiol-Gastr L. 288, G1055–1065.

46. Gout T, Ostor AJ & Nisar MK (2011). Lower gastrointestinal perforation in rheumatoid arthritis patients treated with conventional DMARDs or tocilizumab: a systematic literature review. Clin Rheumatol. 30, 1471–1474.

47. Kuhn KA, Manieri NA, Liu TC & Stappenbeck TS (2014). IL-6 stimulates intestinal epithelial proliferation and repair after injury. PloS one. 9, e114195.

48. Ruhl A, Franzke S, Collins S & Stremmel W (2001). Interleukin-6 expression and regulation in rat enteric glial cells. Am J Physiol-Gastr L. 280, G1163–G1171.

49. Buckley MM, O’halloran KD, Rae MG, Dinan TG & O’malley D (2014). Modulation of enteric neurons by interleukin-6 and corticotropin-releasing factor contributes to visceral hypersensitivity and altered colonic motility in a rat model of irritable bowel syndrome. J Physiol. 592, 5235–5250.

50. Bowen JM, Tsykin A, Stringer AM, Logan RM, Gibson RJ & Keefe DM (2010). Kinetics and regional specificity of irinotecan-induced gene expression in the gastrointestinal tract. Toxicology. 269, 1–12.

51. Wardill HR, Bowen JM, Van Sebille YZ, Secombe KR, Coller JK, Ball IA, Logan RM & Gibson RJ (2016). TLR4-Dependent Claudin-1 Internalization and Secretagogue-Mediated Chloride Secretion Regulate Irinotecan-Induced Diarrhea. Mol Cancer Ther. 15, 2767–2779.

52. Van Sebille YZA, Gibson RJ, Wardill HR, Ball IA, Keefe DMK & Bowen JM (2018). Dacomitinib-induced diarrhea: Targeting chloride secretion with crofelemer. Int J Cancer. 142, 369–380.

