## supplementary data for "TLR4 mediates Il-6 production in irinotecan-induced mucositis"

| ORDINARY ONE -WAY ANOVA OF Il-6 EXPRESSION IN COLON | | | | | |
| --- | --- | --- | --- | --- | --- |
| Alpha | 0.05 | | | | |
| Tukey's multiple comparisons test | Mean Diff. | 95.00% CI of diff. | Significant? | Summary | Adjusted P Value |
| 6 h WT vehicle vs. 72 h WT vehicle | 0.1407 | -0.3916 to 0.6729 | No | ns | 0.9888 |
| 6 h KO vehicle vs. 72 h KO vehicle | 0.1666 | -0.3893 to 0.7225 | No | ns | 0.9770 |
| 6 h WT irinotecan vs. 72 h WT irinotecan | 0.6152 | 0.1077 to 1.123 | Yes | ** | 0.0086 |
| 6 h KO irinotecan vs. 72 h KO irinotecan | -0.8189 | -1.326 to -0.3114 | Yes | *** | 0.0002 |

| ORDINARY ONE-WAY ANOVA FOR Il-6 EXPRESSION IN ILEUM | | | | | |
| --- | --- | --- | --- | --- | --- |
| Alpha | 0.05 | | | | |
| Tukey's multiple comparisons test | Mean Diff. | 95.00% CI of diff. | Significant? | Summary | Adjusted P Value |
| 6h WT vehicle vs. 72 hWT vehicle | 0.6684 | 0.09734 to 1.239 | Yes | * | 0.0122 |
| 6h WT irinotecan vs. 72 h WT irinotecan | 1.604 | 1.033 to 2.175 | Yes | **** | <0.0001 |
| 6h KO vehicle vs. 72 h KO vehicle | 0.4511 | -0.1199 to 1.022 | No | ns | 0.2146 |
| 6h KO irinotecan vs. 72 h KO irinotecan | -0.6059 | -1.177 to -0.03489 | Yes | * | 0.0307 |

| ORDINARY ONE-WAY ANOVA FOR Il-6R EXPRESSION IN ILEUM | | | | | |
|---|---|---|---|---|---|
| Alpha | 0.05 | | | | |
| Tukey's multiple comparisons test | Mean Diff. | 95.00% CI of diff. | Significant? | Summary | Adjusted P Value |
| 6 h WT vehicle vs. 72 h WT vehicle | -0.2683 | -0.7786 to 0.2420 | No | ns | 0.6993 |
| 6 h KO vehicle vs. 72 h KO vehicle | 0.07010 | -0.4402 to 0.5804 | No | ns | 0.9998 |
| 6 h WT irinotecan vs. 72 h WT irinotecan | 0.3250 | -0.1853 to 0.8352 | No | ns | 0.9259 |
| 6 h KO irinotecan vs. 72 h KO irinotecan | 0.02548 | -0.4848 to 0.5358 | No | ns | 0.4721 |

| ORDINARY ONE-WAY ANOVA FOR Il-6R EXPRESSION IN COLON | | | | | |
|---|---|---|---|---|---|
| Alpha | 0.05 | | | | |
| Tukey's multiple comparisons test | Mean Diff. | 95.00% CI of diff. | Significant? | Summary | Adjusted P Value |
| 6 h WT vehicle vs. 72 h WT vehicle | 0.2478 | -0.4317 to 0.9272 | No | ns | 0.9364 |
| 6 h KO vehicle vs. 72 h KO vehicle | -0.6385 | -1.351 to 0.07412 | No | ns | 0.1075 |
| 6 h WT irinotecan vs. 72 h WT irinotecan | 0.04353 | -0.6360 to 0.7230 | No | ns | >0.9999 |
| 6 h KO irinotecan vs. 72 h KO irinotecan | -0.4271 | -1.107 to 0.2524 | No | ns | 0.4872 |

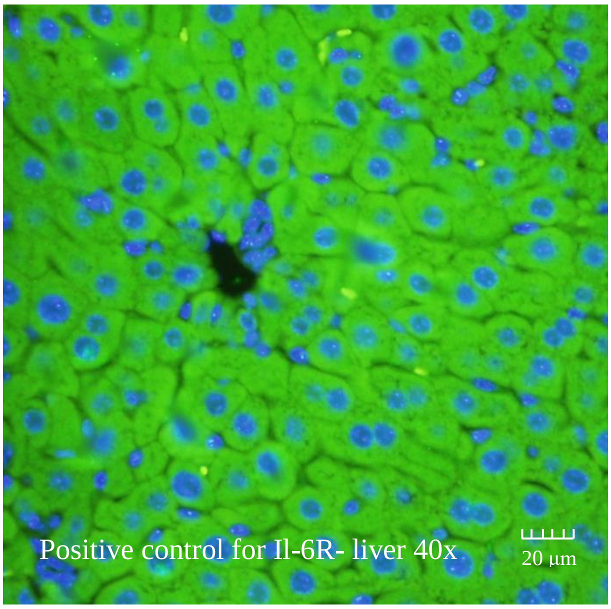
Positive control for Il-6R- liver 40x
20 µm

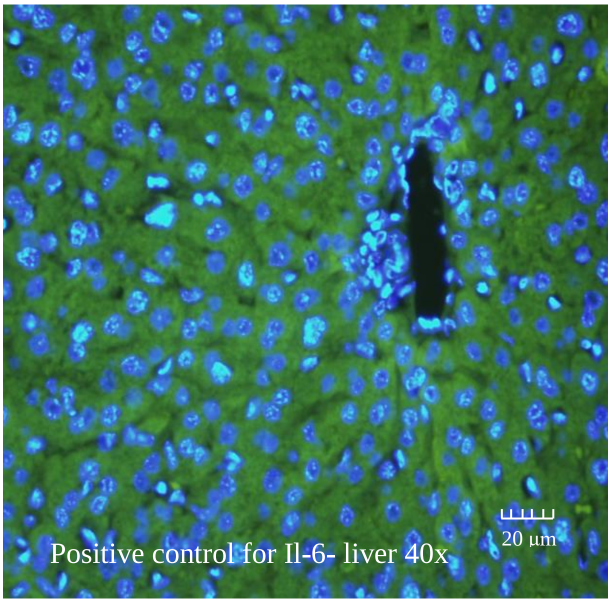
Positive control for Il-6- liver 40x
20 µm

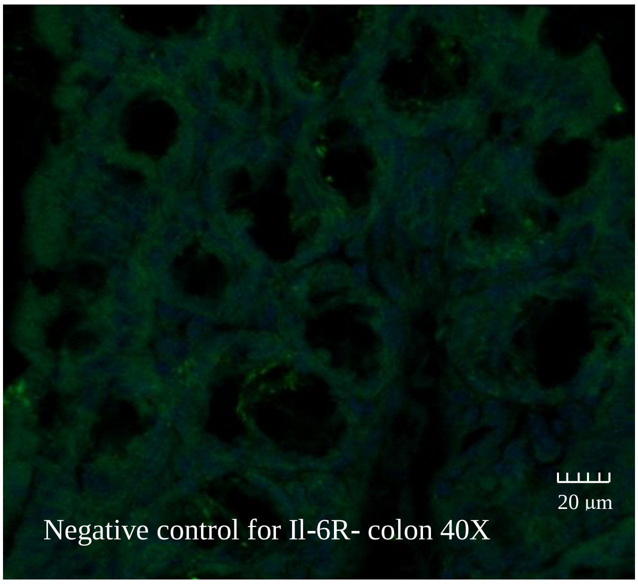
Negative control for Il-6R- colon 40X
20 µm

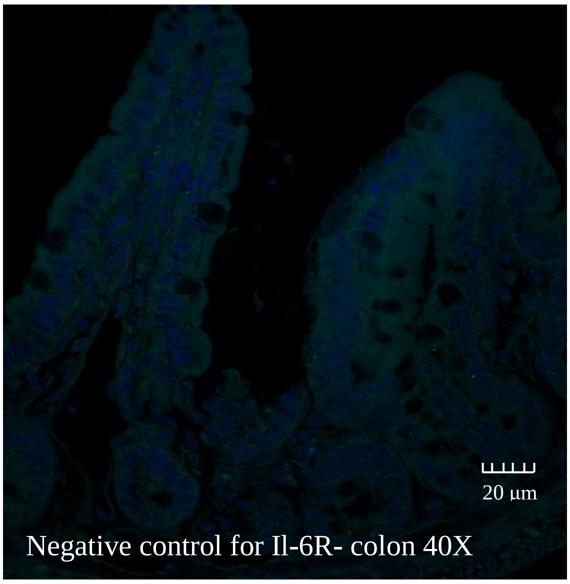
Negative control for Il-6R- colon 40X
20 µm
